# Introducing and benchmarking the accuracy of cayenne: a Python package for stochastic simulations

**DOI:** 10.1101/2020.10.10.334623

**Authors:** Dileep Kishore, Srikiran Chandrasekaran

## Abstract

Biological systems are intrinsically noisy and this noise may determine the qualitative outcome of the system. In the absence of analytical solutions to mathematical models incorporating noise, stochastic simulation algorithms are useful to explore the possible trajectories of these systems. Algorithms used for such stochastic simulations include the Gillespie algorithm and its approximations. In this study we introduce cayenne, an easy to use Python package containing accurate and fast implementations of the Gillespie algorithm (direct method), the tau-leaping algorithm and a tau-adaptive algorithm. We compare the accuracy of cayenne with other stochastic simulation libraries (BioSimulator.jl, GillespieSSA and Tellurium) and find that cayenne offers the best trade-off between accuracy and speed. Additionally, we highlight the importance of performing accuracy tests for stochastic simulation libraries, and hope that it becomes standard practice when developing the same.

The cayenne package can be found at https://github.com/Heuro-labs/cayenne while the bench-marks can be found at https://github.com/Heuro-labs/cayenne-benchmarks

## 1 Introduction

Mathematical modeling and computational simulations play a crucial role in understanding biological systems. Models help simulate the complexity inherent in biological systems, and validate theories and assumptions about various aspects of the system [1]. Biological systems are inherently noisy and hence need mechanical models to incorporate stochasticity in their formulations. Additionally, deterministic models such as ordinary differential equations (ODEs) are unable to capture heterogeneity and complexity in large biological systems [2]. Hence, the reliability of a model greatly depends largely on the chosen modeling approach and specific to the underlying biological problem. ODEs are usually employed when the system under consideration has a large enough number of molecules such that the mean dynamics can be used to represent the behavior of the system and the dynamics of individual molecules can be neglected. Stochastic models on the other hand are used when the molecular populations of the system are lower and where the noise and heterogeneity of the individual molecules or components cannot be neglected [1].

Analytical solutions to stochastic simulations are usually intractable, especially if the system involves a large number of components. The Gillespie algorithm [3, 4] also known as the stochastic simulation algorithm (SSA) or the direct method is a statistically exact method of simulating stochastic models. This method implements a Markov Jump Process (MJP) for simulating a continuous-time Markov chain [3] and is one of the most popular methods by virtue of its algorithmic simplicity. Although the direct method provides exact simulation trajectories, the computational cost of simulating systems with large numbers of entities is very high. This has driven the development of a number of approximate algorithms like the tau-leaping [5] and its improvements (e.g. [6, 7]) making gains in computational speed. These approximate methods take larger steps in time, simulating multiple reactions in each step. Since each individual reaction is not simulated as in the direct method, there is a gain in speed but loss in accuracy. These exact and approximate methods have been implemented in various software libraries; the most notable ones being - BioSimulator.jl [8], GillespieSSA [9], Tellurium [10, 11] and StochKit2 [12]. The open-source scientific computing ecosystem had made Python the de facto standard for leveraging scientific algorithms, but, there is a dearth of open-source Python packages dedicated to stochastic simulations. In this work, we develop the cayenne package - an open-source Python package for performing Gillespie type stochastic simulations. The cayenne package was built with speed, efficiency and ease-of-use in mind and provides a convenient and robust option for simulating small and large biological systems.

A secondary issue addressed in this study pertains to the correctness of implementations of stochastic simulation algorithms in existing software libraries. With the help of the Discrete Stochastic Model Test Suite (DSMTS) [13] we build a benchmarking tool to investigate the accuracy of stochastic simulation algorithms in other libraries, and compare these with cayenne’s performance. We use the results of these benchmarks to identify the algorithms that will serve as good starting points for researchers trying to model their systems stochastically. We also hope that these benchmarks encourage the those developing software for stochastic simulation to incorporate the DSMTS accuracy tests in their libraries.

## 2 Methods

### 2.1 cayenne

The cayenne package is an open-source Python package for performing Gillespie type stochastic simulations. It is built with speed, efficiency and ease-of-use in mind and provides a convenient and robust option for simulating small and large biological systems. The package is easy to install for those familiar with Python, often requiring only a single command through pip. Although the package is written in Python, the speed is not compromised since the algorithms are implemented with Cython rather than native Python. The package also supports the antimony modeling language [10] allowing for the description of the models through text in a simple syntax and avoid the need to specify stoichiometric matrices or rate equations. The underlying propensity functions are restricted to those that are discussed in [3], namely reactions of orders 0-3. Hence, processes such Michaelis-Menten kinetics, which do not fall in the category above, need to be described at the level of their elementary reactions. The code for the package is open-source and can be found at https://github.com/Heuro-labs/cayenne.

The following code section describes the process of simulating a model using the cayenne package. The model `dsmts-003-01` is simulated using the direct algorithm for 10 repetitions.

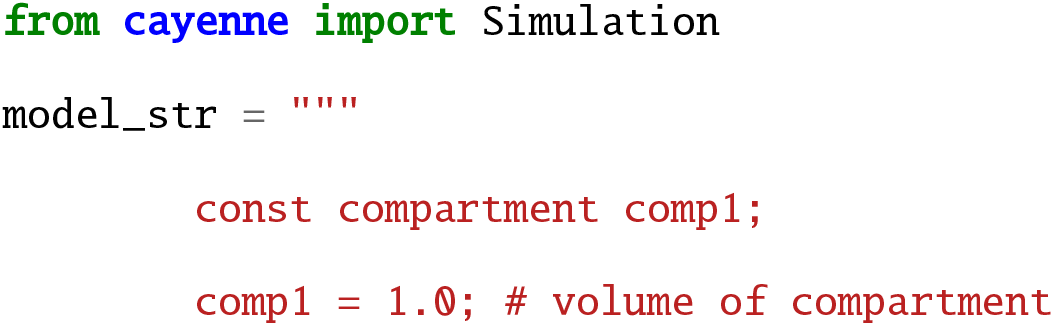
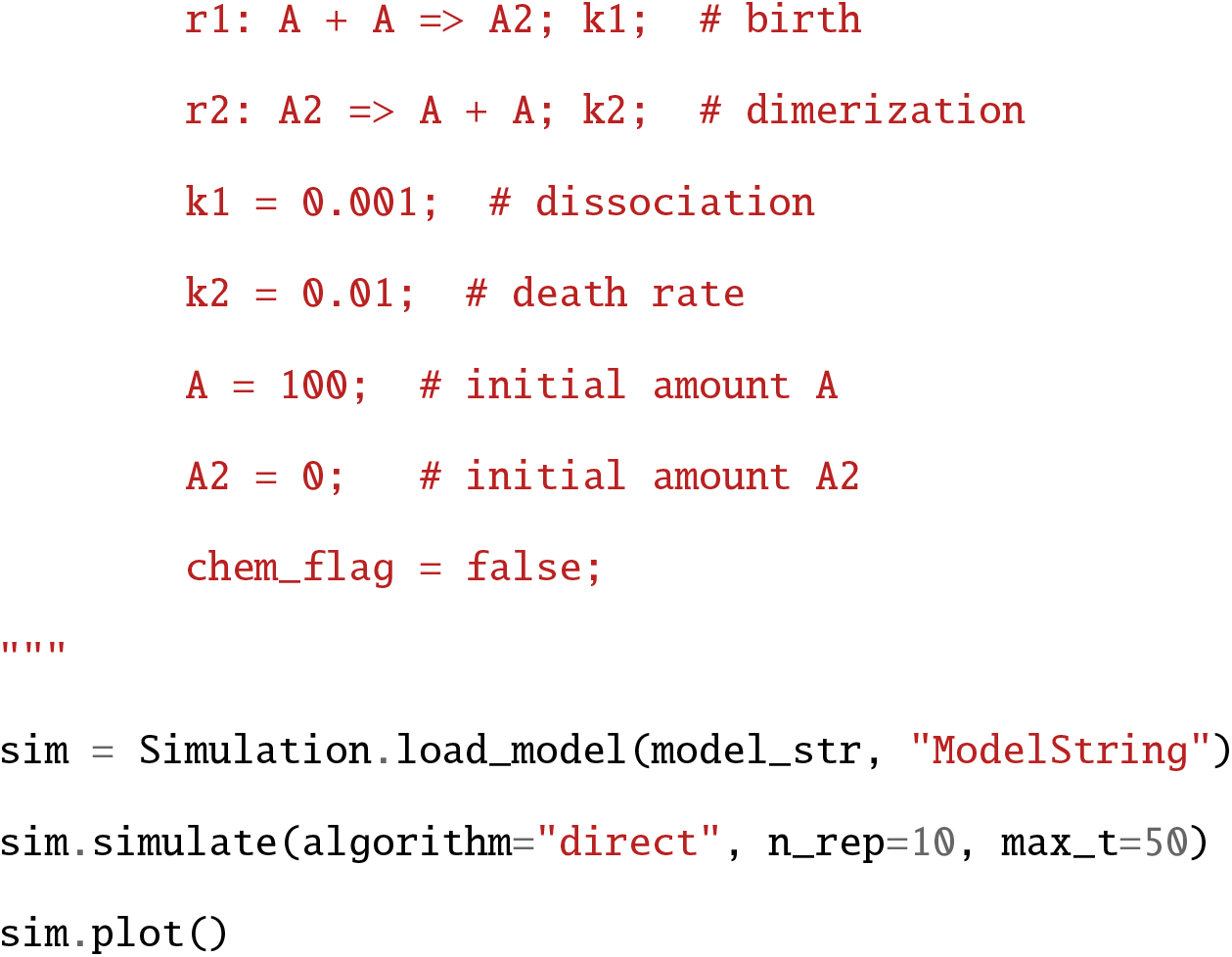

### 2.2 Models

The accuracy of the algorithms in our package were determined using the SBML Discrete Stochastic Model Test Suite (DSMTS) [13]. This test suite presents four reaction systems (each with several parameter combinations) and is intended to verify the accuracy of an exact stochastic simulator that reads models written in the Systems Biology Markup Language (SBML) format [1]. For each of these models (model = system + parameter combination), the distributions of the mean and standard deviation are known analytically. Performing a large number of simulations with the simulator and comparing the distributions of the outputs with these analytical results helps evaluate the simulator’s accuracy.

In addition to verifying the accuracy of the simulator, DSMTS also has tests to verify functionality associated with Systems Biology Markup Language (SBML). For example, model `dsmts-001-02` is identical to `dsmts-001-01`, but the rate parameters are declared to be local instead of global. Since cayenne does not support all features of SBML, such as global and local rate parameters, we only select the unique models (system + parameter combination) for testing. The complete list of models is provided in Table 1.

**Table 1:**
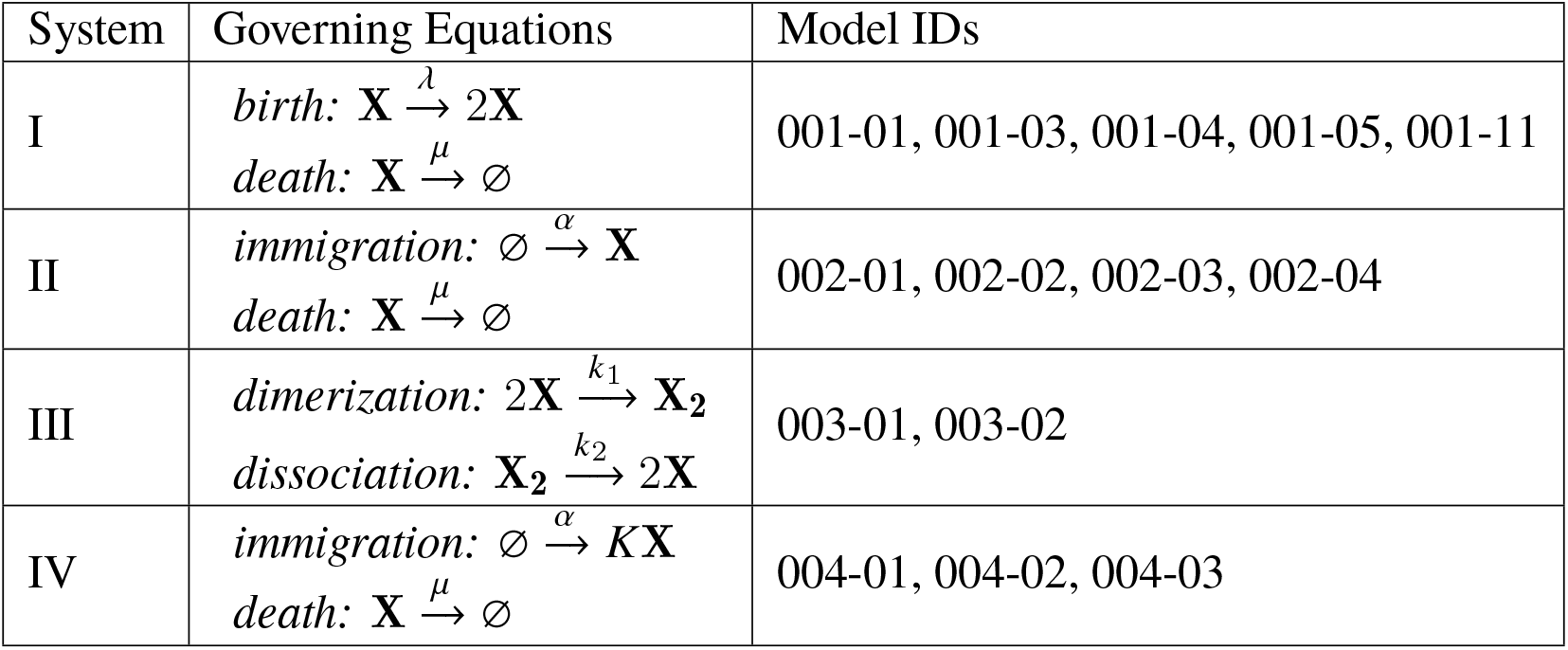
Systems, their equations and the associated models

| System | Governing Equations | Model IDs |
| --- | --- | --- |
| I | $\text{birth: } \mathbf{X} \xrightarrow{\lambda} 2\mathbf{X}$<br>$\text{death: } \mathbf{X} \xrightarrow{\mu} \emptyset$ | 001-01, 001-03, 001-04, 001-05, 001-11 |
| II | $\text{immigration: } \emptyset \xrightarrow{\alpha} \mathbf{X}$<br>$\text{death: } \mathbf{X} \xrightarrow{\mu} \emptyset$ | 002-01, 002-02, 002-03, 002-04 |
| III | $\text{dimerization: } 2\mathbf{X} \xrightarrow{k_1} \mathbf{X}_2$<br>$\text{dissociation: } \mathbf{X}_2 \xrightarrow{k_2} 2\mathbf{X}$ | 003-01, 003-02 |
| IV | $\text{immigration: } \emptyset \xrightarrow{\alpha} K\mathbf{X}$<br>$\text{death: } \mathbf{X} \xrightarrow{\mu} \emptyset$ | 004-01, 004-02, 004-03 |

### 2.3 Libraries and algorithms compared

To contextualize the performance of cayenne with that of other simulation software, we compared the accuracy and speed of cayenne with the following stochastic simulation libraries: Tellurium v2.1.5 [10] (Python v3.6.9), GillespieSSA v0.6.1 [9] (R v3.6.1) and BioSimulator.jl [8] v0.9.3 (Julia v1.3.0).

From these libraries, we assembled algorithms under three labels: (i) direct, (ii) tau-leaping and (iii) adaptive tau-leaping. Of these, the direct algorithms are exact simulators based on the solution to the chemical master equation, based on Gillespie’s early work [3]. The tau-leaping algorithms are approximations. They use a fixed time step over the course of the simulation, firing multiple reactions in each time step. The implementations of this algorithm in all libraries are based on Gillespie’s publication in 2001 [5]. For the tau-leaping simulations in this study we use a fixed tau of 0.03, as it was a good assumption for the models under testing.

For the adaptive tau algorithms, the actual algorithms implemented depended on the library. While cayenne and GillespieSSA use the algorithm from [6] whereas for BioSimulator.jl, we picked the implementation of the algorithm from [7].

### 2.4 Accuracy

#### 2.4.1 Accuracy test for exact simulators

For each model, DSMTS provides the analytical mean (*μ*_*t*_) and standard deviations (*σ*_*t*_) at *t* = 0, 1, .., 50. For exact simulators, the observed mean at time 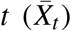 is compared with the analytical mean using the test statistic *Z*_*t*_ defined below [13]:

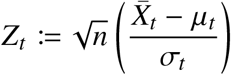

This test is declared as passed if *Z*_*t*_ ~ *N* (0, 1), or if *X*_*t*_ ∈ (−3, 3).

The observed standard deviation at time *t* is compared with the observed standard deviation at time 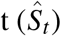 with the test statistic *Z*_*t*_ defined below [13]:

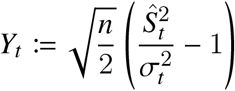

The simulator passes the test if *Y*_*t*_ ~ *N* (0, 1). Since this test is approximate, this test is declared as passed if *Y*_*t*_ ∈ (−5, 5).

#### 2.4.2 Accuracy test for approximate simulators

For the approximate simulators, we used a different test as recommended in the DSMTS user guide. For means, we check if 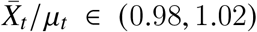. For standard deviations, we check if 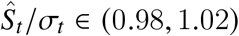.

#### 2.4.3 Accuracy score

To facilitate easy comparison, we defined an accuracy score for each simulation of the model carried out using a particular algorithm implementation. This accuracy score is the percentage of mean and standard deviation tests passed by that algorithm implementation for that model. For example, if an algorithm from a library passes *n*_1_ out of 50 mean tests and *n*_2_ out of 50 standard deviation tests (one test for each time point at which we have analytical values in DSMTS), accuracy = *n*_1_ + *n*_2_. For models where analytical values are known for two species, there are 100 mean tests (50 time points * 2 species) and 100 standard deviation tests. In this case, passing *n*_1_ mean tests and *n*_2_ standard deviation tests gives an accuracy of 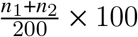. For each combination of model and algorithm implementation, we repeated the simulation 10,000 times (with different random seeds) to calculate the means and standard deviations at each time point. Figure 1 shows the accuracy scores for the model `dsmts-001-003` simulated using the direct algorithm implementation from cayenne.

**Figure 1:**
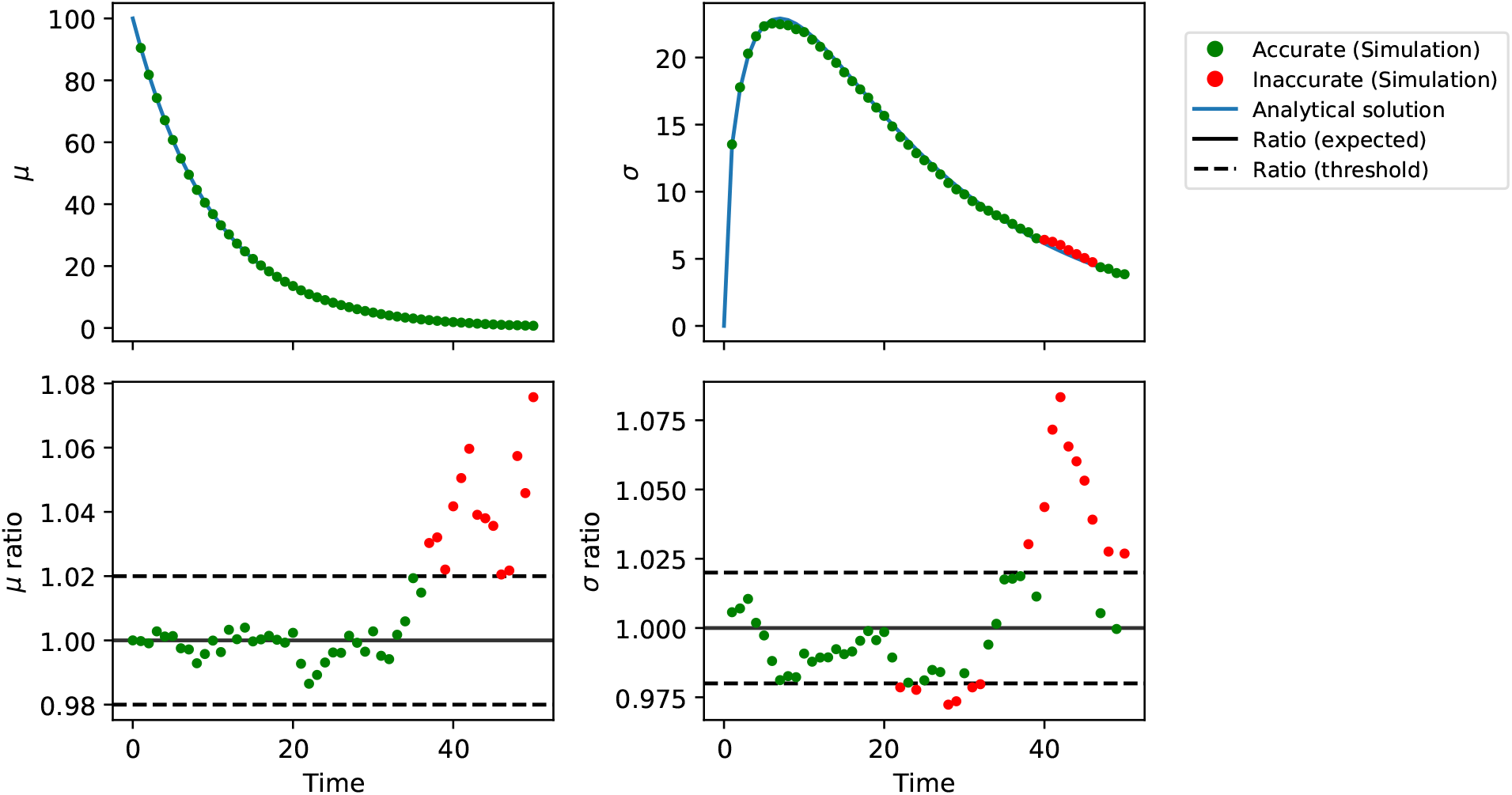
Evaluating the accuracy of a stochastic simulation. Model `dsmts-001-03` is simulated for 10,000 repetitions using the direct algorithm implementation from cayenne. The mean (A) and standard deviation (B) are shown, where the blue lines represent the analytical values and the dots represent the simulated values. The ratio of the means (C) and standard deviations (D) determine the accuracy of the simulation when an approximate algorithm is used. The green dots indicate that the values fall within the acceptable range, whereas the red dots indicate that the values are outside the acceptable range.

### 2.5 Speed evaluation for all simulators

For the speed benchmarks, we picked one model from each of the systems with the exception of the first system, where we picked two models (`dsmts-001-01` and `dsmts-001-03`). For each of these models and algorithm implementations, we ran 10,000 repetitions and measured the time taken. This process was repeated a total of 7 times to estimate the mean time taken. While running the accuracy tests, we found that bugs in some algorithm implementations caused erroneous termination of the simulation instantaneously. Combined with the poor accuracy of other simulations, we believe that a visualization of algorithm run time for a model is best understood in the context of its accuracy score. Hence we present the results of the speed-accuracy comparison in Fig. 6.

### 2.6 Interpolation

For performing the accuracy tests, we require the outputs of the stochastic solvers at integer time points (i.e, *t* = 1, 2, …50). However, running solvers like the direct method and the adaptive tau methods once will output species quantities at non-equispaced time-points (e.g, *t* = 0.17, 0.23, 0.29). Further, running multiple repetitions will generate values at time points that are unique to each repetition. This presents the challenge of aligning solver outputs to integer time points, for example to calculate the mean and standard deviation at each time point required for the accuracy scores. In a more general case, quantities might be needed at non-integral time points, which presents the same problem.

From the theory of the direct method, it is known that the value returned by the solver at time *t*_*i*_ is maintained until it changes at the next time point returned by the solver *t*_*i*+1_ [3]. That is, the value for any time point *t* lying in the interval [*t*_*i*_, *t*_*i*+1_) is simply the value *t*_*i*_. cayenne uses this to extract the value of each repetition at the required integer time points for direct method implementations. Mean and standard deviations are easily computed from the aligned time series.

One potential solution for the tau-leaping algorithm is to pick a tau that is a factor of the time-points we are interested in. For example, for integer time points, we may pick a tau = 0.2 instead of 0.3. This partially alleviates the problem. However, if the tau choice was not suitable for some required intermediate time-point, simulations will have to be repeated which is computationally expensive. Additionally, this problem still persists for methods with adaptive tau algorithms, where time points from different repetitions will not line-up. In these cases, we used a linear interpolation to estimate values for each repetition. The lower and upper bound of the interpolation, for time = *t*_*i*+1_, are simply the species values at *t* and *t*_*i*_ such that *t* ∈ [*t*_*i*_, *t*_*i*+1_).

The BioSimulator package provided two choices of output time points. The first is the raw output of the algorithm, in which time points are not aligned across repetitions. The second provides outputs at requested time points, which are aligned by default. In our study, we included the latter (labeled BioSimulator). We also included the former (unaligned time points) interpolated with cayenne’s interpolation scheme described above (labeled BioSimulator-CI, for cayenne interpolation). This proved to have major effects on the accuracy score and is described in detail in Figure 3.

## 3 Results

### 3.1 cayenne

The cayenne package was as a Python package for Gillespie type stochastic simulations. The output plot generated by the snippet of code in the Methods (Section 2.1) is presented in Figure 2. The code snippet performs a simulation of the system defined by the model `dsmts-003-01` using the cayenne package and plots the result.

**Figure 2:**
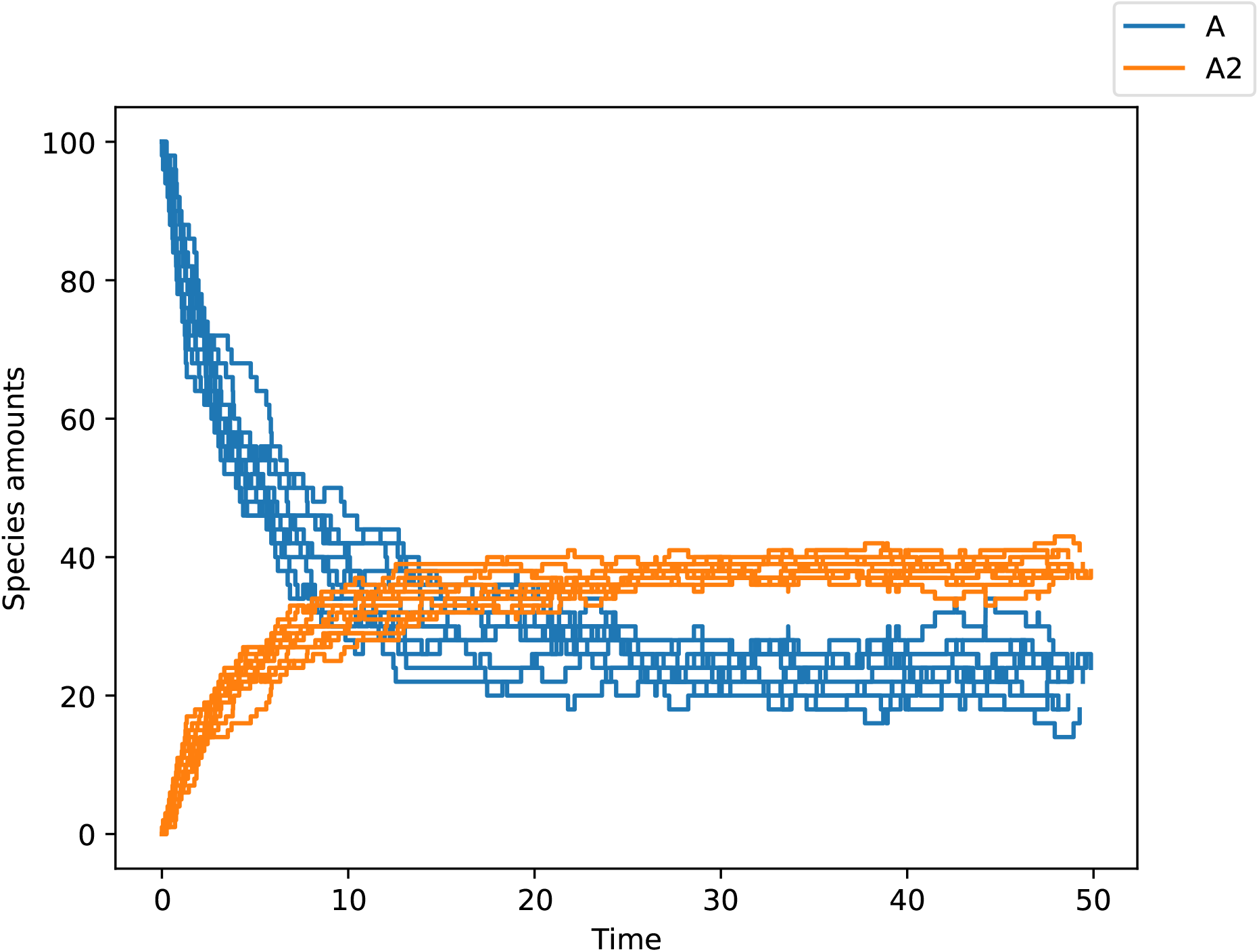
Simulation of model dsmts-003-01 for 10 repetitions using the direct method from the cayenne package. The initial amounts of A (blue) and A2 (orange) were set to 100 and 0 respectively and the simulation was carried out for time *t* = 50.

We have organized the results section as follows. Placing focus on the algorithm, we present the accuracy scores for all models and libraries for a given algorithm. En-route, we also note any algorithm specific or model specific behaviors. Next we discuss the speed of the different algorithms in relation to their accuracy.

### 3.2 Accuracy

The direct algorithm is accurate across software libraries and models (Figure 3). In models `dsmts-001-05` and `dsmts-002-04`, the species amounts increase exponentially, and the direct algorithm did not finish because of the enforced code runtime cutoff. Tellurium performs better than other packages for these models (`dsmts-001-05` and `dsmts-002-04`), but this is expected given that Tellurium’s implementation is not identical to Gillespie’s direct method. Another point of note is the performance drop for model `dsmts-001-03`, which has been observed elsewhere and noted in the DSMTS repository.

**Figure 3:**
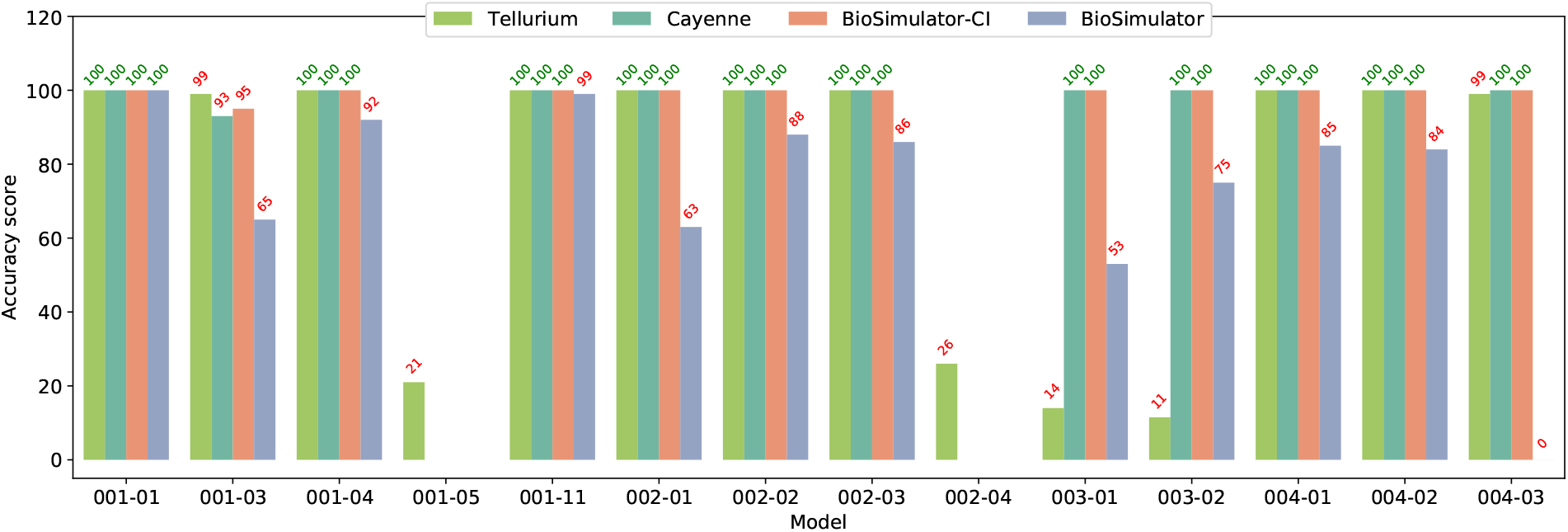
Comparison of the accuracy of stochastic simulations using the direct algorithm implementations. The bar plot shows the accuracy score for each model simulated using the different direct algorithm implementations. Cayenne, BioSimulator-CI and Tellurium are accurate for most of the models, only failing in models `dsmts-001-05` and `dsmts-002-04` because the simulations failed to finish on time. Tellurium does not pass some tests for second order models (`dsmts-003-01` and `dsmts-003-02`) because of possible inaccuracies in its handling of second-order models. GillespieSSA was not used for the comparison because runtimes of the simulations for the direct implementation were very high.

**Figure 4:**
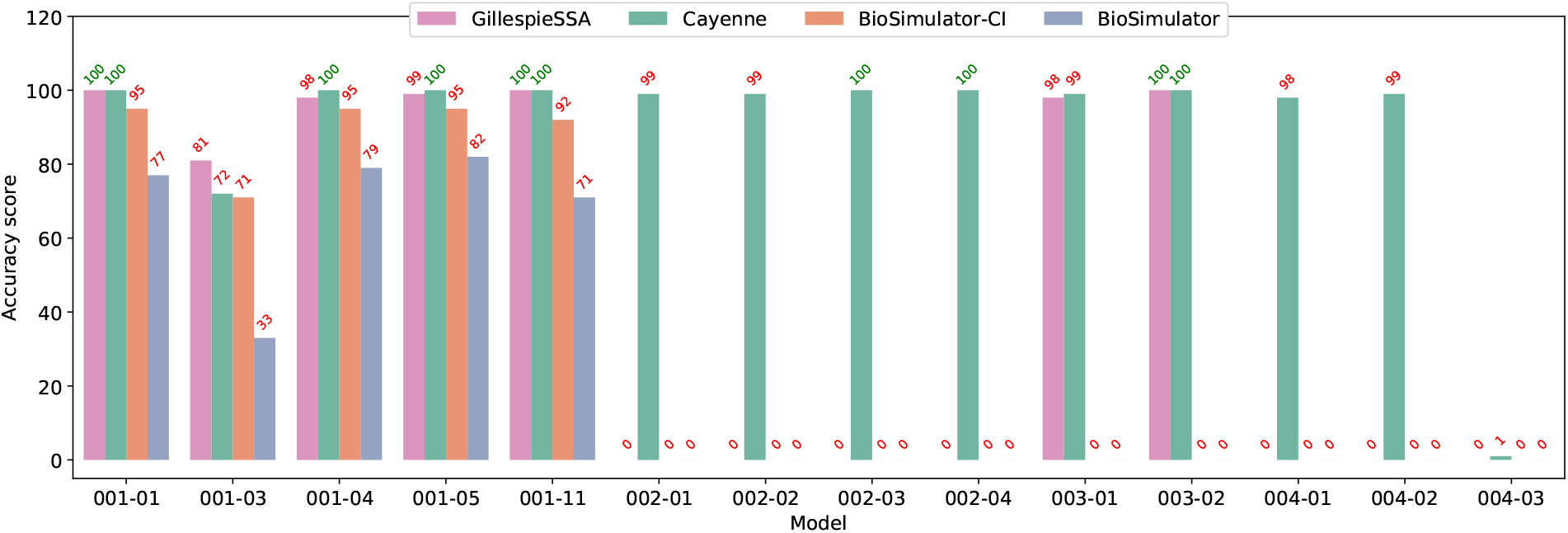
Comparison of the accuracy of stochastic simulations using tau-leaping algorithm implementations. The bar plot shows the accuracy score for each model simulated using the different tau-leaping algorithm implementations. Cayenne is accurate for most of the models, only failing for model `dsmts-004-03` which is a stiff system and does not perform well under approximate algorithms. Incomplete exit-conditions in the tau-leaping implementations are the cause of the failures of BioSimulator* (exits when any of the initial species is 0) and GillespieSSA (exits when all of the initial species are 0) for most of the models. Tellurium was not used for the comparison because it does not have a tau-leaping implementation.

**Figure 5:**
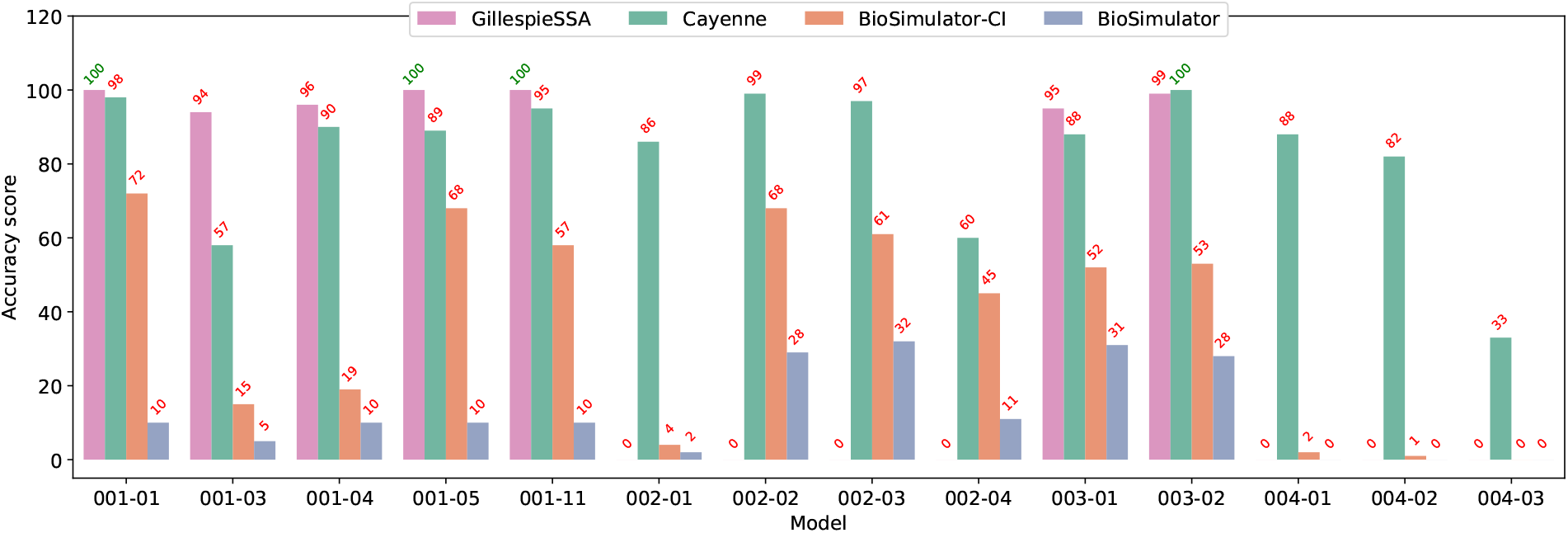
Comparison of the accuracy of stochastic simulations using tau-adaptive algorithm implementations. The bar plot shows the accuracy score for each model simulated using the different tau-adaptive algorithm implementations. GillespieSSA is more accurate than Cayenne and BioSimulator* for most of the models. Incomplete exit-conditions in the tau-adaptive implementations are the cause of the failures of GillespieSSA (exits when all of the initial species are 0) for most of the models. Tellurium was not used for the comparison because it does not have a tau-adaptive implementation.

An interesting observation is the difference in performance of BioSimulator-CI when compared with BioSimulator. Cayenne’s interpolation combined with BioSimulator.jl’s raw simulation output provides better accuracy. This suggests that the internal implementation of the direct algorithm is correct in the BioSimulator.jl package, but that the interpolation step used there leads to some inaccuracies. A closer investigation of the error plots for these cases suggests that it is in areas of high slope that this difference in interpolation becomes important (Figure S1, Figure S2).

We also noticed that Tellurium is not accurate for models `dsmts-003-01` and `dsmts-003-02`. These are part of System III, and one point of difference between this system and others is that System III is a second order system. Therefore it is possible that Tellurium’s implementation of the second and possibly higher order systems is inaccurate.

The performance for the tau-leaping algorithm in general is poorer than the direct algorithm. This is to be expected as the tau-leaping algorithm is approximate while the direct is exact. This does not explain the large gaps in performance we see, especially for GillespieSSA and BioSimulator.jl. This is because the tau-leaping implementations in these packages do not cover some corner cases; i) GillespieSSA does not begin simulation for any model where all of the initial species amounts are 0 ii) BioSimulator.jl does not begin simulation for any model where any of the initial species amounts are 0.

In cases not affected by these bugs, GillespieSSA and cayenne do equally well. BioSimulator-CI is slightly behind when combined with cayenne’s implementation. BioSimulator (with BioSimulator.jl’s native interpolation) appears to consistently fall short without cayenne’s interpolation, showing that care must be taken when converting raw simulation results into a structure expected by the end-user (such as values at specific time-points of interest). Tau-leaping performs worse in model `dsmts-001-03` than the direct algorithm, likely because it is an approximation technique. Tau-leaping is however able to solve models `dsmts-001-05` and `dsmts-002-04` in the allotted time with good accuracy, since it makes constant time step leaps and does not attempt to simulate every event in an exponentially growing system.

The adaptive tau algorithms largely perform as well as the tau-leaping algorithms. GillespieSSA’s implementation still carries the bug seen in the tau-leaping algorithm (simulation exits when all initial species are 0). Interestingly, BioSimulator doesn’t carry over the bug seen in its tau-leaping implementation. Where not affected by bugs, GillespieSSA’s adaptive-tau algorithm performs better than cayenne’s implementation. Both perform better than BioSimulator with or without cayenne’s interpolation.

### 3.3 Speed

We do not present a comparison of algorithm speeds agnostic of their accuracy, since bugs in the implementation can deflate run times to a large extent. However, we do perform an accuracy vs. speed comparison for a set of models (Figure 6).

**Figure 6:**
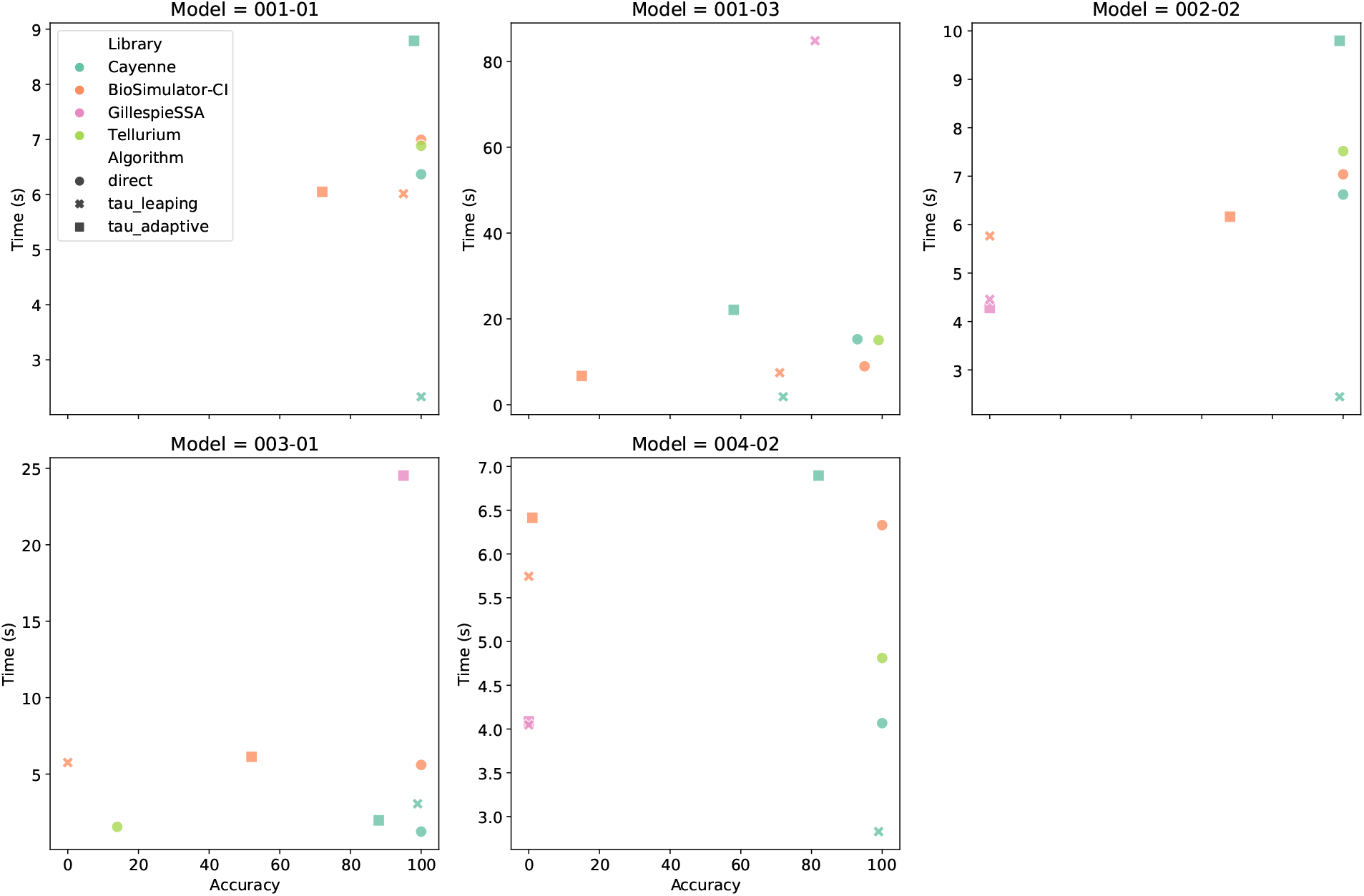
Accuracy and speed comparison of libraries and models. Each point represents the accuracy score (X axis) and mean run time (Y axis) of a given algorithm and library combination. Points within a given panel belong to the same model.

When contextualized with speeds, there was significant variability between the libraries and algorithms across the different models as shown in Figure 6. We focus our observations on the cases where accuracy score was at least 80. The tau-leaping algorithm was faster than the direct algorithm for 4 out of 5 models across all libraries, but the opposite is true for model `dsmts-003-01` in which the direct algorithm was the fastest. Interestingly, cayenne’s tau-adaptive was also the slowest for 4 out of 5 models with model `dsmts-003-01` being the exception. We also observe that GillespieSSA is slower than the other libraries, likely due to differences in speed between native R and compiled code in Python/Julia. Other trends in speed appear to be model specific, suggesting that the model structure and parameters determine the order of library and algorithm performance.

## 4 Discussion

In this study, we developed the cayenne package for performing Gillespie type stochastic simulations. The package is easy to install and use for those familiar with Python. Yet, the speed is not compromised since the algorithms are implemented with Cython rather than native Python. Using a subset of the antimony modeling language, we also provide users with a method to quickly prototype their models, alleviating the need to specify stoichiometric matrices or rate equations.

We compared the performance of the algorithms in our package with other existing packages, uncovering trends that would be useful for audiences who develop models (modelers) as well as those who develop software packages and stochastic simulation algorithms used by the modelers.

From the perspective of modelers, we see that both the choice of simulation algorithm and the choice of the library implementing that algorithm are important. We interpret our results to recommend the following choice of algorithms for modelers. First, Gillespie’s direct algorithm should be tried, along with a tested interpolation scheme for reliable outputs at specified time-points.

Should this algorithm be too slow for the intended purposes, a naïve implementation of tau-leaping can be pursued. The naïve tau-leaping algorithm is the building block for several adaptive methods. Consequentially, it is one of the simplest algorithms and easiest to implement correctly. While a value of *τ* = 0.03 worked well for the models tested in this study, a modeler may need to tune their value of *τ*. A simple heuristic to identify a good *τ* is checking if the results do not change significantly when *τ* is reduced (e.g., halved). Models which are stiff (e.g. model `dsmts-004-03`) may require a lower *τ*, or the choice of an adaptive algorithm. From our accuracy studies, we see that adaptive tau algorithms score lower than tau-leaping counterparts - across different libraries considered. Moreover, the adaptive tau algorithms are not always faster than naïve tau-leaping. Modelers who choose to use the adaptive tau algorithms should be cognizant of the significant drops in accuracy seen in simple models.

Additionally, the choice of library appears to matter as well. For example, barring bugs seen in the 002 and 004 series of models, GillespieSSA’s implementation of the adaptive-tau algorithm of Cao et. al 2006 [6] is more accurate than cayenne’s. The differences observed across different algorithms, and especially across packages, call for more rigorous standards in library and algorithm development. We hope that the results of our study encourage package developers and those who invent algorithms to check the accuracy performance of algorithms before releasing them. Adopting the DSMTS test-suite would ensure a fair comparison among different methods. They will also serve as a reference for future implementations of the algorithm, ensuring greater reproducibility in the long run.

While the DSMTS test-suite is a good standard for verifying accuracy of stochastic simulation algorithms, there are some key model properties that are not represented in its repertoire. For example, none of the models exhibit oscillations. Arguably, something of greater importance is stochastic bifurcation [14]. An example of this: when simulated long enough, the result of a competition between two equally capable bacterial species will result in the survival of one species and the extinction of the other. However, the mean of all trajectories will suggest survival of both species. Systems of this kind, where no actual trajectory resembles the mean of all trajectories, are currently not represented in the DSMTS test-suite. Hence, the findings of this study may not be applicable for such models, and other cases not represented by the DSMTS test-suite.

An interesting outcome of the study is the consistently better performance of BioSimulator-CI in comparison with BioSimulator. This suggests that careful interpolation of simulation trajectories are important for achieving high accuracy of estimates at specified time points. This difference is stark even for the direct algorithm, where BioSimulator-CI achieves perfect scores in several cases missed by BioSimulator. Hence although the underlying algorithm is correct, reported values at specified time points are inaccurate. Linear interpolation for the other algorithms also improves results, but does not result in full scores. The cayenne package thus offers a good trade-off between accuracy and speed, compared to BioSimulator (less accurate, with some bugs) and GillespieSSA (some bugs, slower). Of course, simple bugs may be fixed in future versions of these libraries. This will enable a more rigorous accuracy comparison across the packages.

## Supplementary

**Figure S1:**
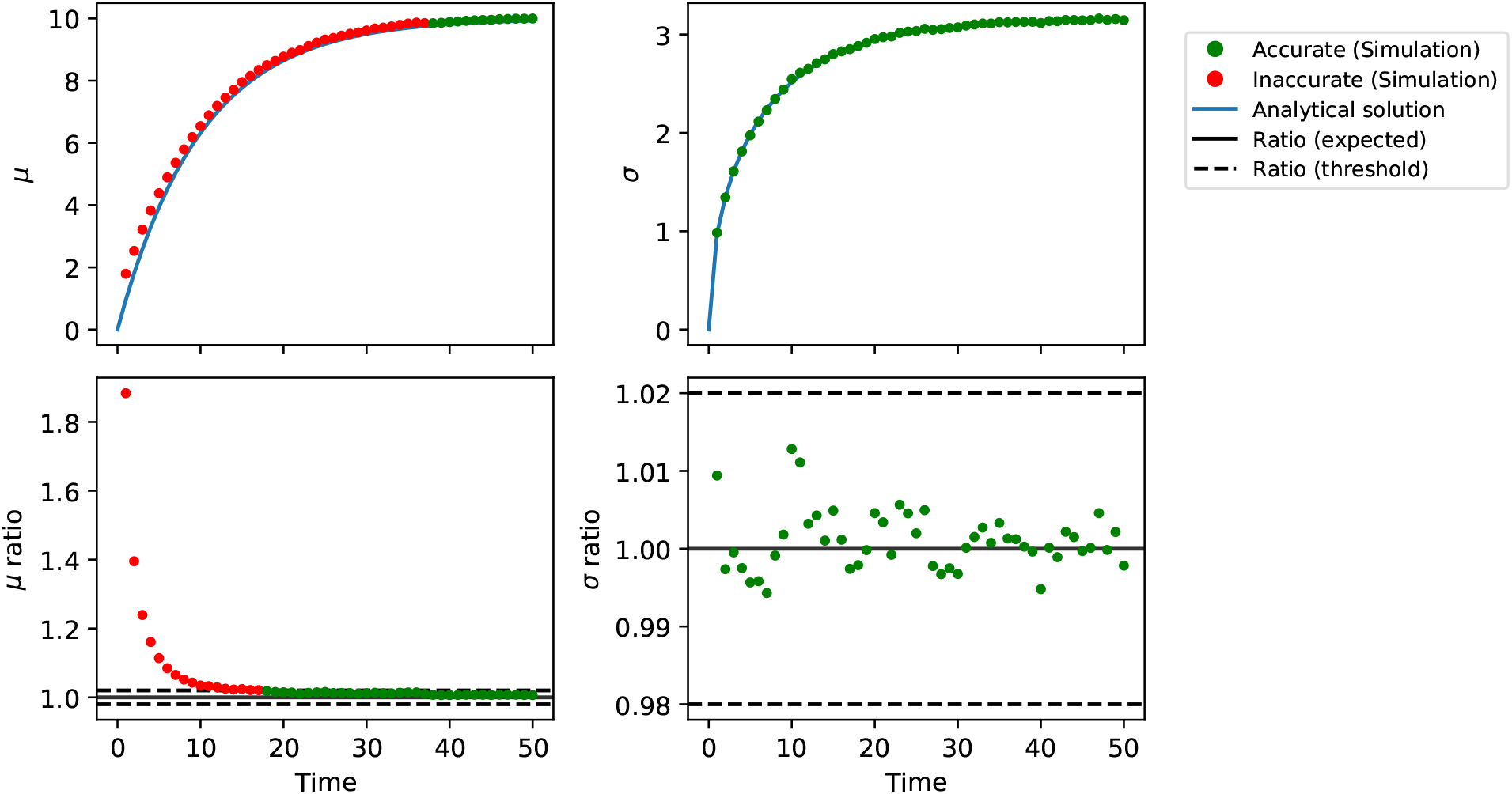
Evaluating the accuracy of the simulation of model dsmts-002-01 using the BioSimulator direct method implementation. Model `dsmts-002-01` is simulated for 10,000 repetitions using the direct algorithm implementation from BioSimulator. The mean (A) and standard deviation (B) are shown, where the blue lines represent the analytical values and the dots represent the simulated values. The ratio of the means (C) and standard deviations (D) determine the accuracy of the simulation when an approximate algorithm is used. The green dots indicate that the values fall within the acceptable range, whereas the red dots indicate that the values are outside the acceptable range.

**Figure S2:**
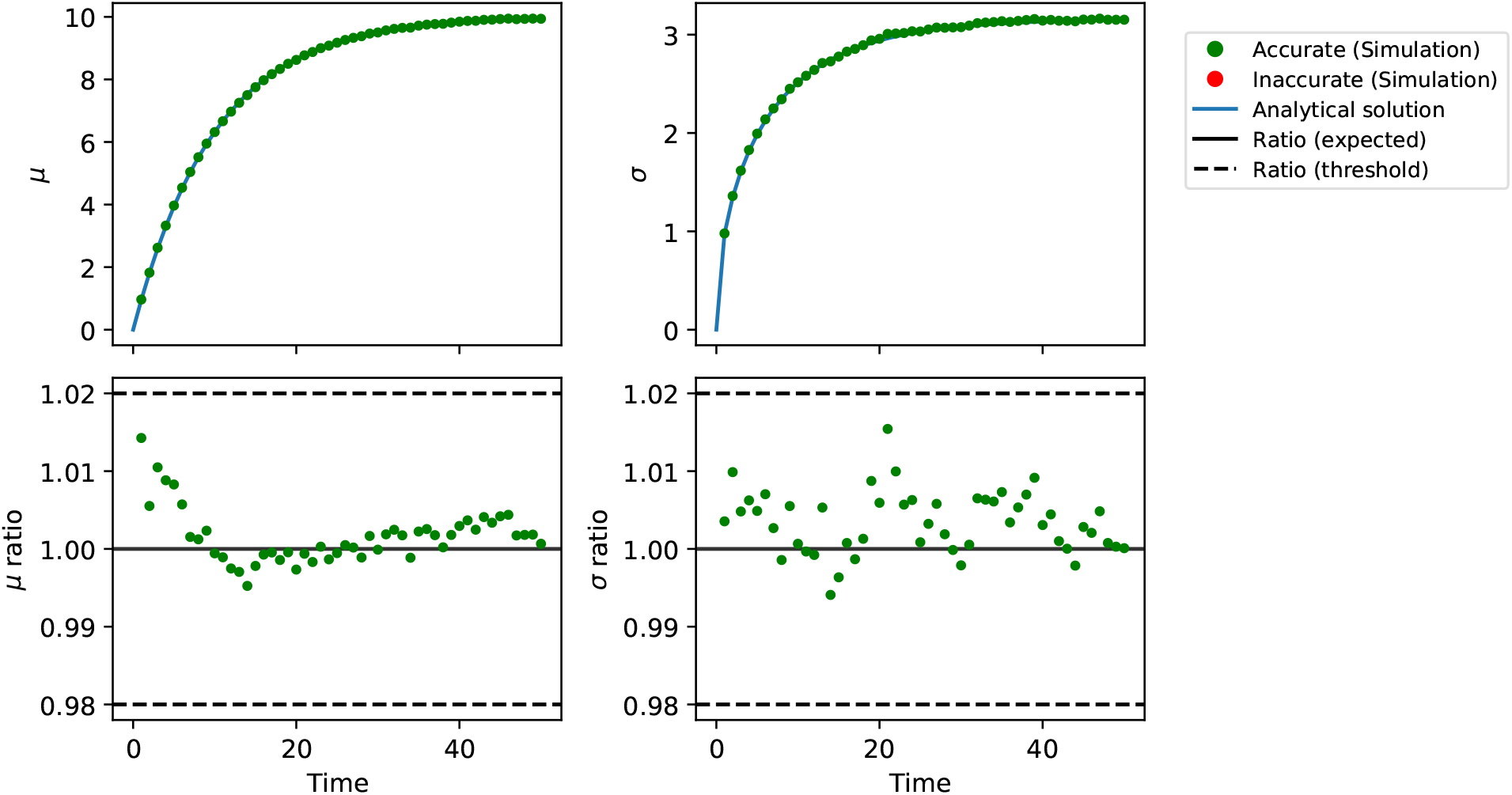
Evaluating the accuracy of the simulation of model dsmts-002-01 using the BioSimulator-CI direct method implementation. Model `dsmts-002-01` is simulated for 10,000 repetitions using the direct algorithm implementation from BioSimulator with cayenne’s interpolation (BioSimulator-CI). The mean (A) and standard deviation (B) are shown, where the blue lines represent the analytical values and the dots represent the simulated values. The ratio of the means (C) and standard deviations (D) determine the accuracy of the simulation when an approximate algorithm is used. The green dots indicate that the values fall within the acceptable range, whereas the red dots indicate that the values are outside the acceptable range.

